# High-throughput phenogenotyping *of Mycobacteria tuberculosis* clinical strains reveals bacterial determinants of treatment outcomes

**DOI:** 10.1101/2023.04.09.536166

**Authors:** Sydney Stanley, Caitlin N. Spaulding, Qingyun Liu, Michael R. Chase, Dang Thi Minh Ha, Phan Vuong Khac Thai, Nguyen Huu Lan, Do Dang Anh Thu, Nguyen Le Quang, Jessica Brown, Nathan D. Hicks, Xin Wang, Maximillian Marin, Nicole C. Howard, Andrew J. Vickers, Wiktor M. Karpinski, Michael C. Chao, Maha R. Farhat, Maxine Caws, Sarah J. Dunstan, Nguyen Thuy Thuong Thuong, Sarah M. Fortune

## Abstract

**Background:** Combatting the tuberculosis (TB) epidemic caused by *Mycobacterium tuberculosis* (*Mtb*) necessitates a better understanding of the factors contributing to patient clinical outcomes and transmission. While host and environmental factors have been evaluated, the impact of *Mtb* genetic background and phenotypic diversity is underexplored. Previous work has made associations between *Mtb* genetic lineages and some clinical and epidemiological features, but the bacterial traits underlying these connections are largely unknown.

**Methods:** We developed a high-throughput functional genomics platform for defining genotype-phenotype relationships across a panel of *Mtb* clinical isolates. These phenotypic fitness profiles function as intermediate traits which can be linked to *Mtb* genetic variants and associated with clinical and epidemiological outcomes. We applied this approach to a collection of 158 *Mtb* strains from a study of *Mtb* transmission in Ho Chi Minh City, Vietnam. *Mtb* strains were genetically tagged in multiplicate, which allowed us to pool the strains and assess *in vitro* competitive fitness using deep sequencing across a set of 14 host-relevant antibiotic and metabolic conditions. Phylogenetic and monogenic associations with these intermediate traits were identified and then associated with clinical outcomes.

**Findings:** *Mtb* clinical strains have a broad range of growth and drug response dynamics that can be clustered by their phylogenetic relationships. We identified novel monogenic associations with *Mtb* fitness in various metabolic and antibiotic conditions. Among these, we find that mutations in *Rv1339*, a phosphodiesterase, which were identified through their association with slow growth in glycerol, are further associated with treatment failure. We also identify a previously uncharacterized subclade of Lineage 1 strains (L1.1.1.1) that is phenotypically distinguished by slow growth under most antibiotic and metabolic stress conditions *in vitro*. This clade is associated with cavitary disease, treatment failure, and demonstrates increased transmission potential.

**Interpretation:** High-throughput phenogenotyping of Mtb clinical strains enabled bacterial intermediate trait identification that can provide a mechanistic link between *Mtb* genetic variation and patient clinical outcomes. *Mtb* strains associated with cavitary disease, treatment failure, and transmission potential display intermediate phenotypes distinguished by slow growth under various antibiotic and metabolic conditions. These data suggest that Mtb growth regulation is an adaptive advantage for host bacterial success in human populations, in at least some circumstances. These data further suggest markers for the underlying bacterial processes that govern these clinical outcomes.

**Funding:** National Institutes of Allergy and Infectious Diseases: P01 AI132130 (SS, SMF); P01 AI143575 (XW, SMF); U19 AI142793 (QL, SMF); 5T32AI132120-03 (SS); 5T32AI132120-04 (SS); 5T32AI049928-17 (SS) Wellcome Trust Fellowship in Public Health and Tropical Medicine: 097124/Z/11/Z (NTTT) National Health and Medical Research Council (NHMRC)/A*STAR joint call: APP1056689 (SJD) The funding sources had no involvement in study methodology, data collection, analysis, and interpretation nor in the writing or submission of the manuscript.

**Research in context:** *Evidence before this study:* We used different combinations of the words mycobacterium tuberculosis, tuberculosis, clinical strains, intermediate phenotypes, genetic barcoding, phenogenomics, cavitary disease, treatment failure, and transmission to search the PubMed database for all studies published up until January 20^th^, 2022. We only considered English language publications, which biases our search. Previous work linking *Mtb* determinants to clinical or epidemiological data has made associations between bacterial lineage, or less frequently, genetic polymorphisms to *in vitro* or *in vivo* models of pathogenesis, transmission, and clinical outcomes such as cavitary disease, treatment failure, delayed culture conversion, and severity. Many of these studies focus on the global pandemic Lineage 2 and Lineage 4 *Mtb* strains due in part to a deletion in a polyketide synthase implicated in host-pathogen interactions. There are a number of *Mtb* GWAS studies that have led to novel genetic determinants of *in vitro* drug resistance and tolerance. Previous *Mtb* GWAS analyses with clinical outcomes did not experimentally test any predicted phenotypes of the clinical strains. Published laboratory-based studies of *Mtb* clinical strains involve relatively small numbers of strains, do not identify the genetic basis of relevant phenotypes, or link findings to the corresponding clinical outcomes. There are two recent studies of other pathogens that describe phenogenomic analyses. One study of 331 *M. abscessus* clinical strains performed one-by-one phenotyping to identify bacterial features associated with clearance of infection and another details a competition experiment utilizing three barcoded *Plasmodium falciparum* clinical isolates to assay antimalarial fitness and resistance.

*Added value of this study:* We developed a functional genomics platform to perform high-throughput phenotyping of *Mtb* clinical strains. We then used these phenotypes as intermediate traits to identify novel bacterial genetic features associated with clinical outcomes. We leveraged this platform with a sample of 158 *Mtb* clinical strains from a cross sectional study of *Mtb* transmission in Ho Chi Minh City, Vietnam. To enable high-throughput phenotyping of large numbers of *Mtb* clinical isolates, we applied a DNA barcoding approach that has not been previously utilized for the high-throughput analysis of *Mtb* clinical strains. This approach allowed us to perform pooled competitive fitness assays, tracking strain fitness using deep sequencing. We measured the replicative fitness of the clinical strains in multiplicate under 14 metabolic and antibiotic stress condition. To our knowledge, this is the largest phenotypic screen of *Mtb* clinical isolates to date. We performed bacterial GWAS to delineate the *Mtb* genetic variants associated with each fitness phenotype, identifying monogenic associations with several conditions. We then defined *Mtb* phenotypic and genetic features associated with clinical outcomes. We find that a subclade of *Mtb* strains, defined by variants largely involved in fatty acid metabolic pathways, share a universal slow growth phenotype that is associated with cavitary disease, treatment failure and increased transmission potential in Vietnam. We also find that mutations in *Rv1339*, a poorly characterized phosphodiesterase, also associate with slow growth *in vitro* and with treatment failure in patients.

*Implications of all the available evidence:* Phenogenomic profiling demonstrates that *Mtb* strains exhibit distinct growth characteristics under metabolic and antibiotic stress conditions. These fitness profiles can serve as intermediate traits for GWAS and association with clinical outcomes. Intermediate phenotyping allows us to examine potential processes by which bacterial strain differences contribute to clinical outcomes. Our study identifies clinical strains with slow growth phenotypes under *in vitro* models of antibiotic and host-like metabolic conditions that are associated with adverse clinical outcomes. It is possible that the bacterial intermediate phenotypes we identified are directly related to the mechanisms of these outcomes, or they may serve as markers for the causal yet unidentified bacterial determinants. Via the intermediate phenotyping, we also discovered a surprising diversity in *Mtb* responses to the new anti-mycobacterial drugs that target central metabolic processes, which will be important in considering roll-out of these new agents. Our study and others that have identified *Mtb* determinants of TB clinical and epidemiological phenotypes should inform efforts to improve diagnostics and drug regimen design.

## Introduction

*Mycobacterium tuberculosis* (*Mtb*) is responsible for over 10 million cases of tuberculosis (TB) disease and 1·5 million deaths per year.^1^ Mitigating the global burden of disease is challenging because the determinants of TB treatment success, disease outcomes, and transmission dynamics involve a combination of host, environmental, and pathogen factors.^2–4^ In particular, we have a relatively superficial understanding of the bacterial determinants of clinically relevant outcomes beyond high-level drug resistance, although these discoveries are actionable where the genetic basis of the relevant phenotypes is defined.^5, 6^

Recent studies have sought to use the wealth of *Mtb* whole genome sequencing data to find novel genetic associations with drug resistance, several of which have been experimentally shown to cause altered drug responses.^7–10^ A few studies have sought to find bacterial genetic associations with more complex clinical outcomes like treatment failure, transmission, and dissemination.^10–16^ Interpretation of genetic associations is complicated by *Mtb*’s high degree of genetic linkage.^17^ In addition, the biologic basis of most clinical phenotypes and the functions of most *Mtb* genes *in vivo* are unclear, and combinations of genomic changes may also work in concert to produce the overall clinical phenotype of interest. Therefore, the biologic plausibility of any given association is often uncertain.

The best studied putative determinant of *Mtb* infection outcome is a deletion in a gene encoding a polyketide synthase (*pks15/1*), which synthesizes phenolic glycolipid (PGL), an immunologically active cell wall lipid.^18^ This deletion was originally identified in *Mtb* Lineage 4 (L4) strains via a targeted comparison with strains from the epidemiologically expanding clade of Lineage 2 (L2.2), and the intact gene is proposed to account for the population level fitness success of the L2.2 subfamily.^19, 20^ Subsequent work demonstrated that this was an overly simplistic model as PGL production can be found across a number of *Mtb* lineages and indeed is the ancestral rather than the evolutionarily derived state.^21, 22^

The gap between genetics and complex patient phenotypes is not a problem unique to TB. The mechanistic study of many human diseases faces the same challenge. As a potential solution, human geneticists turned to intermediate trait analysis to dissect the genetic determinants of disease and their mechanistic contributions.^23^ Intermediate phenotypes are heritable biological traits shaped by the same genetic drivers as the more complicated disease processes under investigation. The intermediate phenotypes may be related to the clinical phenotype of interest in non-intuitive ways. However, they can still provide a genetic handle for, and mechanistic insights into, otherwise intractable complex clinical states. For example, identifying cognitive variables and deficits as intermediate phenotypes of mechanistically impenetrable disorders such as schizophrenia and depression has aided in the identification and interpretation of genes associated with these diseases.^23^

In this study, we developed a high-throughput phenogenomic platform for the discovery of *Mtb*-intrinsic intermediate traits related to two clinical phenotypes: treatment failure and development of lung cavitary disease. Cavitary disease is an indicator of TB disease severity and is associated with treatment failure, while both outcomes are linked to transmission.^24, 25^ By bridging the gap between *Mtb* genetic diversity and patient outcomes via bacterial intermediate phenotypes, we show that *Mtb* clinical strains demonstrate a wide range of phenogenomic variation in response to metabolic and drug stressors, and identify novel bacterial genetic and phenotypic correlates with cavitary disease, treatment failure, and potentially transmission.

## Methods

### *Mtb* clinical strains

We sought to analyze a representative subset of 1,635 *Mtb* strains collected through population-based study of *Mtb* transmission dynamics in Ho Chi Minh City, Vietnam (S1 Table).^11^ *Mtb* strains were isolated from the sputum of HIV negative, smear positive adults seeking care at the district TB units from 2008-2011.^11^ We began with a set of 200 strains selected to be representative of the lineage distribution of the cohort. We successfully barcoded 158 strains; attrition occurred due to various factors including failed strain recovery after transport from Vietnam, inability to confirm phenotypic drug sensitivity, and dropout in the barcoding process (unsuccessful transformations, contamination, and overlapping barcodes).

### Patient clinical metadata and ethics

The strains are paired to corresponding high-level patient data including age, sex, lung cavitation identified via chest radiography, and treatment outcome.^12^ Treatment failure was defined as persistent smear positivity after 5 months of appropriate treatment.^13^ The study was approved by the institutional research board of Pham Ngoc Thach Hospital, Ho Chi Minh City, Vietnam and the Oxford University Tropical Research Ethics Committee (OxTREC 030–07). Written informed consent was obtained from all patients for enrolment into the study.

### Drug sensitivity testing

The clinical strains in our study were previously determined to be phenotypically drug sensitive to ethambutol (EMB), isoniazid (INH), rifampicin (RIF), and streptomycin (SM) with the BACTEC MGIT 960 SIRE system (BD, Franklin Lakes, NJ, USA).^13^ Prior to barcoding, we verified the susceptibility to RIF via a pooled liquid and solid media growth inhibition assay (Extended Methods).

### Barcoded *Mtb* clinical strain library

Each clinical isolate in our study plus a *Mtb* reference strain (Erdman) was transformed with an integrating plasmid library carrying randomized 18bp DNA sequence tag that inserts at the L5 phage integration site of the *Mtb* genome.^26^ Our goal was to create three independent barcoded clones for each strain to serve as biological replicates. Due to technical dropout or exclusion because of barcode sequence similarity, 66 strains in the library are represented by three clones, 64 strains by two clones and 29 strains by a single clone. In total, the library contains 355 unique barcodes. The barcoded strains were pooled together in roughly equal abundances according to OD_600_ to form the input library, which we aliquoted into 1 mL stocks. We sequenced the input library to determine representation of each barcode (fig. S1) (Extended Methods).

### *In vitro* competition experiments and data analysis

We inoculated the input library into liquid standard media (7H9), defined single carbon source media, and standard media treated with each antibiotic at the indicated concentrations and 20 μg/mL of kanamycin to maintain selection for the barcode (Extended Methods). Each condition was prepared in triplicate for technical replicates. We measured the OD_600_ of the cultures at day 3 (D3) and day 6 (D6) after inoculation to track bulk library growth (fig. S2). At D3 and D6 we also extracted gDNA from the cultures and the stock input library for deep sequencing to quantify strain barcode abundance. To compare barcode abundance in the technical replicates, we calculated the barcode normalized read counts as following, which controls for differences in sample read depth and barcode abundance in the input library (S2 Table):

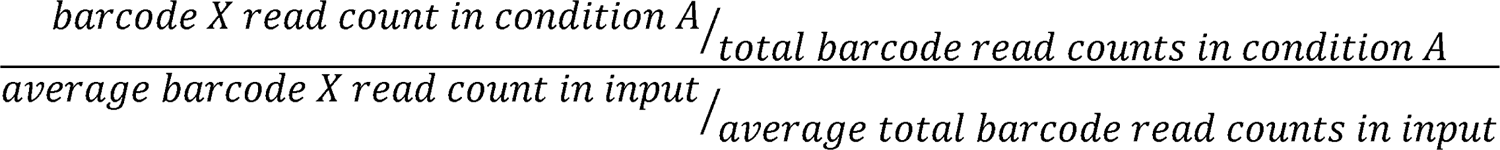

To compare barcode abundance among the biological replicates, identical *Mtb* clinical strains with unique barcodes, we averaged the normalized read counts across the technical replicates for each barcode for each condition (S2 Table). To obtain the relative fitness (RF) values for each strain that serve as our intermediate phenotypes, normalized barcode read counts were averaged across the technical and biological replicates and the log_2_ fold change was determined. We calculated the RF coefficient of variation for use in subsequent analysis where the behavior of individual strains is important (S2 Table) (Extended Method). Competition experiments were repeated for the acetate + propionate, glycerol, lactate, BDQ, CFZ, and SM conditions to further assess the reproducibility of the assay (fig. S3, S3 Table).

### Genomic, phylogenetic, and mutation analyses

We performed whole-genome sequencing (WGS) of the 158 barcoded strains used in our study (200 Mbp Illumina DNA sequencing, SeqCenter, Pittsburgh, PA, USA) to ensure the strains we phenotyped were paired to the correct sequences as compared to those determined by Holt *et al.* and to confirm that the strains did not acquire mutations known to confer drug resistance. One uniquely barcoded isolate was sequenced per strain. Refer to the extended methods for the procedure for alignment, variant calling, and mutation annotation. S1 Table includes the strain accession numbers from this study (accession number PRJNA950969) and those provided by Holt *et al.* (accession number PRJNA355614).^11^ We found discordance between our WGS and the original sequences for 18 strains (S1 Table) and used the corrected sequences for our analyses. We analyzed the variant calling files for each clinical isolate to confirm that the strains in our sample did not contain polymorphisms at the loci associated with EMB, INH, RIF, SM, and quinolone resistance as previously assessed by Holt *et al*.^11^

To identify *Mtb* mutations associated with the relative fitness phenotypes, we used the Python (version 3.8.5) package Pyseer (version 1.3.10) to perform a bacterial GWAS.^27^ We used the linear mixed model option and included only nonsynonymous single nucleotide polymorphisms (SNPs) and mutations within intergenic regions for the analysis. Loci were grouped by genes. We controlled for population structure with SNP-based similarity and distance matrices and by inputting the lineage assignment for each strain (Extended Methods).

### Growth Curves

Selected clinical isolates were cultured in 7H9 media and 20 μg/mL kanamycin to an OD_600_ of ∼0·5, then each strain was diluted in triplicate to an OD_600_ of 0·005 into either 10 mL of fresh 7H9 media or acetate + propionate defined medium with 20 μg/mL kanamycin. Cultures were incubated at 37° C with constant shaking and OD_600_ was measured on days 3, 5, and 7.

### Minimal inhibitory drug concentration

We determined the minimum inhibitory concentration (MIC) for the selected clinical isolates using the indicated concentrations of antibiotics via an alamar blue reduction assay completed as described but without shaking.^8^ After 3 days of incubation with the alamar blue reagent (BioRad, Hercules, CA, USA), the MIC was identified as the lowest concentration of drug that inhibited the reagent color change as determined visually.

### Terminal branch length analysis

Terminal branch lengths for the L1 strains were calculated as the number of SNPs derived from the most recent branching point of each strain. A consensus alignment of all variable sites for the L1 WGS within the complete *Mtb* strain set from the population study was generated to compute pairwise SNP distances of all L1 strain pairs.^11^ Then strain pairs with the smallest SNP distances were identified from the SNP distance matrix of all L1 strains. The terminal branch length of one *Mtb* strain is calculated by the SNP distance from its closest neighbor subtracted by the number of SNPs that were accumulated by its closest neighbor.

### Statistical analyses

Prism 9 (version 9.5.0) for Mac OS was used for statistical analyses unless indicated otherwise.

## Results

### Methodology for high-throughput intermediate phenotyping of *Mtb* clinical isolates

We sought to develop a platform to identify bacterial intermediate traits associated with infection and treatment outcomes. As in human genetic studies, we anticipated using these traits to nominate several novel bacterial determinants of clinically relevant outcomes. Drug resistance is a known bacterial determinant of treatment failure, cavitary disease, and other clinical phenotypes; but there is little known about bacterial factors associated with TB disease in drug-sensitive strains.^24, 28, 29^ Therefore, here we aimed to identify novel traits associated with clinical outcomes by performing high dimensional phenotyping of drug susceptible *Mtb* strains as confirmed by phenotypic and genotypic DSTs.

Our sample set is comprised of *Mtb* strains isolated from 158 TB patients enrolled in a population-based study of *Mtb* transmission in Ho Chi Minh City, Vietnam (Fig. 1A).^11^ TB cavitary disease and treatment failure occurred in 27·8% and 8·9% of the patients whose strains are represented in this sample (Table 1). Lineage-level assignment was significantly associated with treatment outcome; 19% of patients in our sample infected by a L1 strain failed treatment, compared to 5·6% of L2 strains and none of the L4 strains (Chi-square statistic=7·7, P=0·022) (Fig. 1A) (Table 2). Despite the relationship between cavitary disease and treatment failure described in the literature, we observed no significant association between these clinical outcomes in our dataset (Chi-square statistic=3·05, P=0·38).^24^

**Figure 1.**
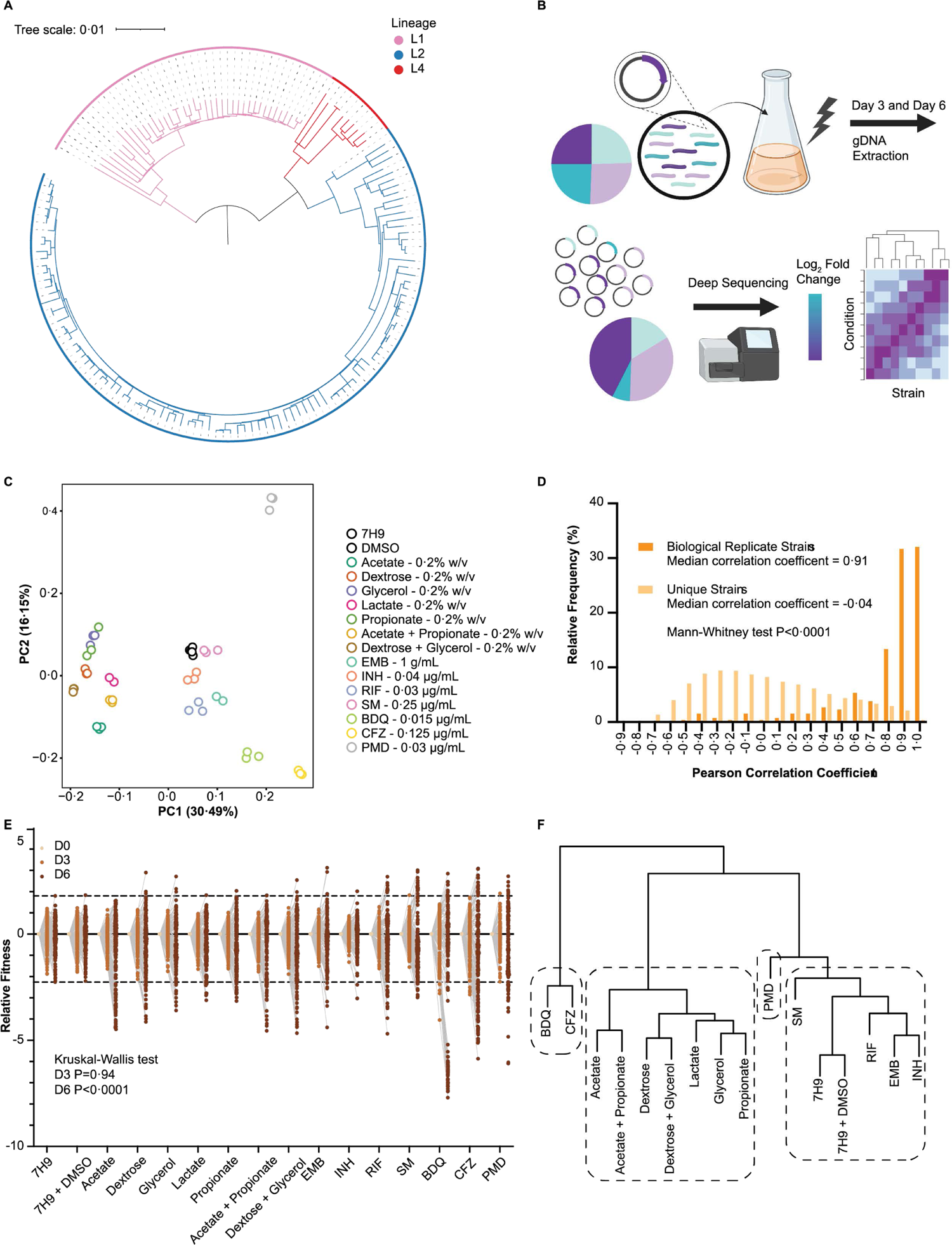
*Mtb* clinical isolates genetically barcoded for pooled competition experiments to examine relative metabolic and antibiotic fitness phenotypes. A. Approximately maximum-likelihood tree of the 158 *Mtb* strains + Erdman selected for the study. The scale indicates the number of mutations per site. Rooted at the midpoint. B. Diagram of the *in vitro* competition experiment workflow using genetically barcoded *Mtb* clinical isolates. Created with BioRender. C. Principal component analysis (PCA) of the technical replicates based on D6 strain normalized barcode read counts. D. For the barcode replicates (isogenic strains with unique barcodes that serve as biological replicates) and the non-replicates (unique strains), histogram shown of Pearson correlation coefficients comparing D6 strain normalized barcode read counts. 159 unique strains with 1-3 different barcodes each, so 355 strains compared in total. E. Dot plot of relative fitness (RF) values for all 159 strains per indicated condition and timepoint. RF values are normalized to strain abundance in input pool inoculum, so all D0 values are set to zero. F. Ward’s linkage clustering of the stress conditions based on strain D6 RF values. Agglomerative coefficient = 0. 76. Boxes denote 4 different k-mean clusters.

**Table 1.** Sample of patients with culture-confirmed tuberculosis disease from Vietnam.

| Demographics of tuberculosis patient sample (n=158) |  |
| --- | --- |
| **Age** |  |
| 15-29 years | 45 (28.5%) |
| 30-49 years | 80 (50.6%) |
| 50-69 years | 30 (19%) |
| 70-89 years | 3 (1.9%) |
| **Sex** |  |
| Male | 114 (72.2%) |
| Female | 44 (27.8 %) |
| **Infecting strain lineage** |  |
| L1 | 42 (26.6%) |
| L2 | 107 (67.7%) |
| L4 | 9 (5.7 %) |
| **Cavitary Disease** |  |
| Yes | 44 (27.8%) |
| No | 114 (72.2%) |
| **Treatment Failure** |  |
| Yes | 14 (8.9%) |
| No | 144 (91.1%) |

**Table 2.** Clinical outcomes associated with culture-confirmed tuberculosis disease from the sample Vietnamese patients.

| Association between demographic features and clinical outcomes within TB patient sample |  |  |  |  |
| --- | --- | --- | --- | --- |
|  | Cavitary disease percentage | Cavitary disease Chi-square | Treatment failure percentage | Treatment failure Chi-square |
| Age, years |  |  |  |  |
| 15-29 | 12/45 (26.7%) | Chi-square = 3.05<br>P=0.38 | 4/45 (8.9%) | Chi-square = 2.46<br>P=0.48 |
| 30-49 | 20/80 (25%) |  | 6/80 (7.5%) |  |
| 50-69 | 10/30 (33.3%) |  | 3/30 (10%) |  |
| 70-89 | 2/3 (66.7%) |  | 1/3 (33.3%) |  |
| Sex |  |  |  |  |
| Male | 32/114 (28.1%) | Chi-square = 0.01<br>P=0.92 | 11/114 (9.6%) | Chi-square = 0.32<br>P=0.57 |
| Female | 12/44 (27.3%) |  | 3/44 (6.8%) |  |
| Infecting Strain Lineage |  |  |  |  |
| L1 | 16/42 (38.1%) | Chi-square = 3.01<br>P=0.22 | 8/42 (19%) | Chi-square = 7.7<br>P=0.022 |
| L2 | 26/107 (24.3%) |  | 6/107 (5.6%) |  |
| L4 | 2/9 (22.2%) |  | 0/9 (0%) |  |

We next sought to better resolve bacterial determinants of clinical outcomes though bacterial intermediate trait identification. To accomplish this, we developed a platform for the high-throughput phenotypic profiling of *Mtb* strains. For this approach we tagged *Mtb* clinical isolates with unique genetic barcodes, enabling us to pool strains and track individual strain abundance using deep sequencing in large-scale high-throughput *in vitro* competition experiments across different host- and clinically relevant stress conditions (Fig. 1B).^26^ From these data, we defined host-relevant metabolic and antibiotic stress fitness phenotypes for each clinical isolate, which is indicative of the relative change of strain relative abundance throughout the time course of the competition experiments.

During infection, *Mtb* sources of carbon is limited by the availability within the host environment.^30^ We therefore utilized defined single carbon source media (acetate, propionate, acetate + propionate, dextrose, glycerol, dextrose + glycerol, and lactate) in order to mimic host-relevant metabolic stress.^31^ Our phenotyping screen also included first-line antibiotics (isoniazid (INH), rifampin (RIF), ethambutol (EMB) and streptomycin (SM)) to which these drug-sensitive isolates may have been exposed but not acquired high level resistance, and second-line antibiotics (bedaquiline (BDQ), clofazimine (CFZ), and pretomanid (PMD)) to which these strains were not expected to have been exposed but which were utilized to probe essential bacterial processes.^32^ We screened the library at antibiotic concentrations at roughly the MIC_50_, the concentration that inhibits growth of 50% of tested clinical isolates on average, to create growth restriction without complete killing.^33–36^

We assessed the technical and biological reproducibility of our phenotyping method. We performed principal component analysis (PCA) of the technical replicates under each condition using the normalized barcode read counts at day 6 (D6), which were normalized to control for differences in read depth and strain abundance in the input library (fig. S1). The clustering of the technical replicates from each condition demonstrates the technical reproducibility of the measurements (Fig. 1C, S2 Table). Biological replicates were achieved by independently constructing up to three barcoded clones per clinical isolate. The median Pearson correlation coefficient of normalized barcode read counts across all conditions is 0·91 when comparing biological replicates of the same strain but is −0·04 for comparisons between different strains (Mann-Whitney P<0·0001, Fig. 1D, S2 Table). Thus, biological replicates are concordant while the lack of correlation between unique strains demonstrates that the clinical isolates in our library are phenotypically diverse. For the biological replicates, any barcode that resulted in a Pearson correlation coefficient of 0·62 or less was considered an outlier via the ROUT test, excluding 32 of 355 barcodes from further analysis (Fig. 1D, S2 Table).^37^ Given the quality of the technical and biological replicates, for each strain we averaged the normalized barcode read counts across replicates and calculated the log_2_ of these values to obtain the corresponding relative fitness (RF) for each condition (S2 Table), which then established the metabolic and antibiotic *Mtb* clinical strain intermediate phenotypes used for downstream association studies. The Spearman correlation coefficients of the RF values from fully independent repeats of the competition experiments ranged from 0·72 to 0·97 (fig. S3, S2 Table, S3 Table).

We observed a large range of fitness phenotypes across the metabolic and antibiotic stress conditions, the distributions of which were significantly different by D6 (Kruskal-Wallis P<0·0001) (Fig. 1E). Hierarchical clustering of the conditions based on strain D6 RF values found similar carbon source conditions (such as acetate and acetate + propionate) and drugs with related mechanisms of action (EMB and INH or BDQ and CFZ) group together (Fig. 1F). The carbon source conditions formed a distinct cluster from the antibiotic conditions (Fig. 1F). Interestingly, PMD forms an outgroup from other drug conditions while BDQ and CFZ form an outgroup from all conditions, indicating the *Mtb* clinical strains have distinct phenotypes in the face of these drugs which target different aspects of energy metabolism (Fig. 1F).

We confirmed that our RF values track with more traditional drug sensitivity phenotypes by performing MIC assays with BDQ, CFZ, PMD, and SM. Strains with the highest RF values demonstrate MICs as much as 15 times greater than that of strains with the lowest RF values, although all were still under the critical concentration defining high level drug resistance (fig. S4, S4 Table).^38^ We also performed single-strain growth curves in acetate + propionate defined media and found that strains with the largest RF values have a faster growth rate compared to strains with the lowest RF values (fig. S5, S4 Table).

### Phylogenetic structure of metabolic and antibiotic intermediate phenotypes

Given the association between L1 *Mtb* clinical strains and treatment failure in our sample (Table 2), we compared differences in the intermediate phenotypes based on lineage. L1 strains exhibit significantly lower RF values compared to strains belonging to L2 and L4 across nearly every stress condition (Fig. 2). This phenotype is especially striking in acetate, acetate + propionate, dextrose + glycerol, BDQ, and CFZ (Fig. 2). A notable exception is PMD, where L1 strains demonstrate higher RF values compared to L2 and L4 strains (Fig. 2).

**Figure 2.**
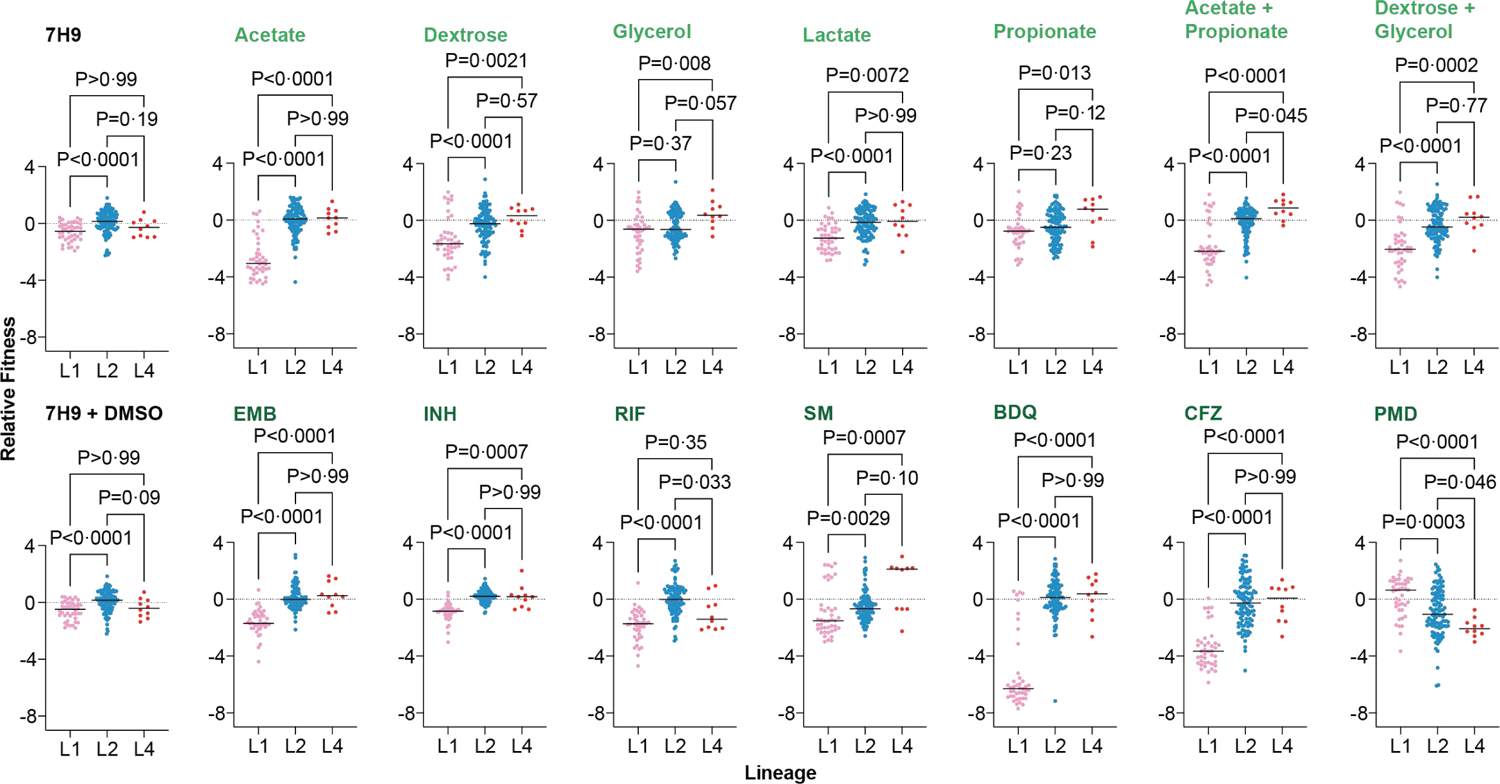
Lineage patterns of *Mtb* clinical isolates antibiotic and metabolic relative fitness phenotypes. Dot plot of strain RF values for the indicated conditions, grouped by lineage. Kruskal-Wallis p-values adjusted by the Dunn’s multiple comparisons test indicated.

To assess the genetic basis of the RF phenotypes, we organized the RF values according to the phylogenetic relationship of the clinical isolates (Fig. 3A). Within-lineage phenotypic heterogeneity is evident but lineage and sub-lineage patterning of the stress response phenotypes is qualitatively recognizable. For each condition we determined the phylogenetic signal, which quantifies the propensity of closely related strains to share similar intermediate traits compared to more genetically distant isolates.^39^ We found that the RF of strains under every condition except lactate, EMB, and PMD is associated with a phylogenetic signal of 0·5 or higher (P<0·0001) which indicates that most phenotypes have high phylogenetic heritability (fig. S6).

**Figure 3.**
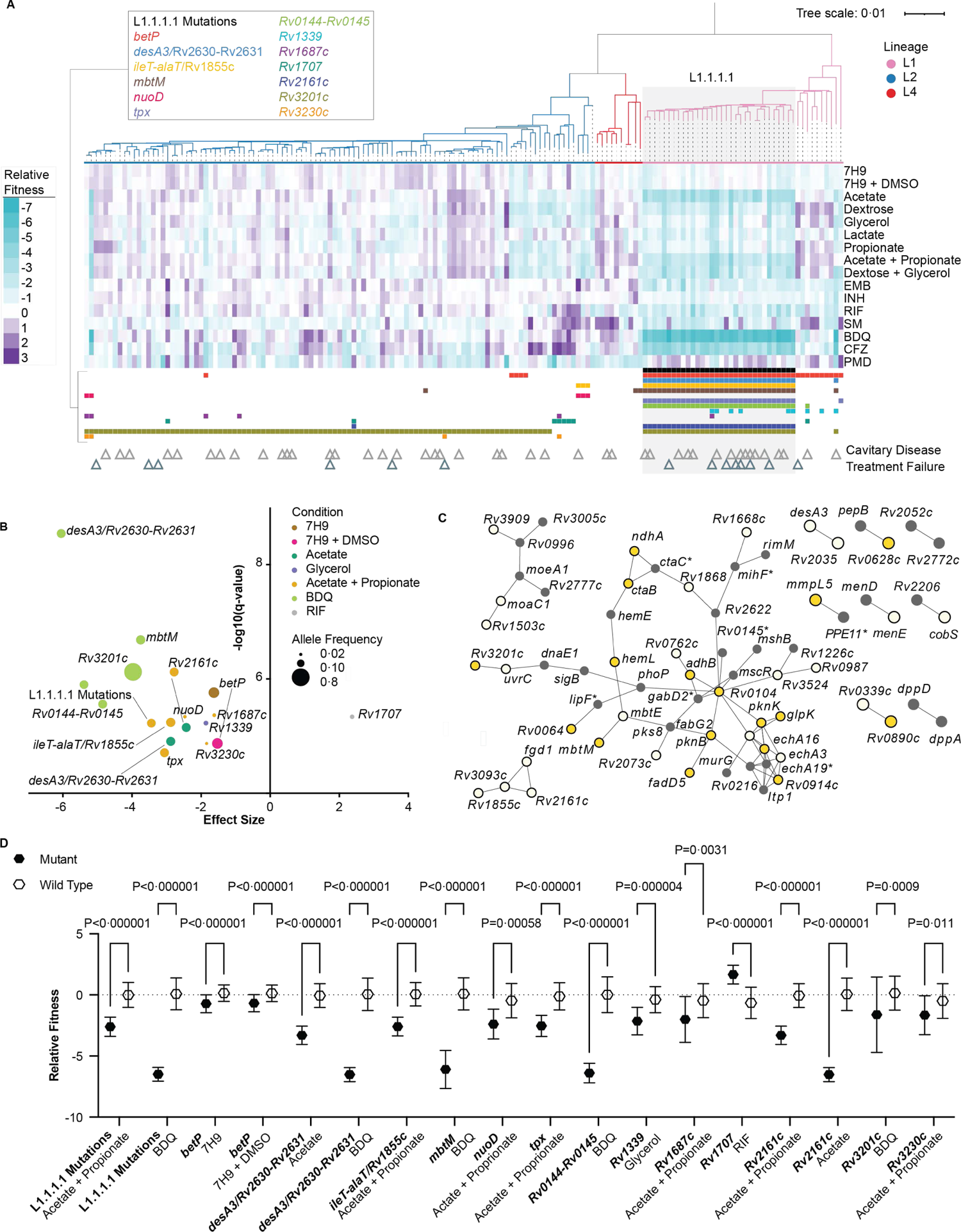
*Mtb* clinical strain relative fitness phenotypes are heterogeneous but follow genetic and sub-lineage associated patterns. A. Heatmap of D6 RF values ordered according to the Fig. 1 phylogeny. Triangles indicate clinical phenotypes associated with each strain. Color squares mark strains carrying mutations in the indicated genes returned as significant hits from the GWAS. B. Bubble plot of the significant hits from the GWAS. Allele frequency refers to the proportion of strains within the clinical isolate sample carrying at least one mutation in the indicated gene. C. STRING network analysis of the genes carrying mutations unique to L1.1.1.1. Intolerant nonsynonymous mutations predicted by SIFT or premature termination mutations (dark yellow node); tolerant and indeterminate nonsynonymous mutations (light yellow node); synonymous mutations (black node); intergenic mutations (black node and asterisk). For intergenic mutations, the nearest gene downstream of the mutation is indicated. D. Dot plots comparing relative fitness values of mutant and wild type strains in the indicated conditions for the significant gene hits from the GWAS. Dots indicate mean, error bars indicate standard deviation. P-value results of Mann-Whitney test shown after Benjamini, Krieger, and Yekutieli multiple test correction indicated.

### *Mtb* genetic determinants underlying intermediate phenotypes

We sought to explicitly define genetic determinants of the intermediate phenotypes using a GWAS. We performed GWAS using a linear mixed model approach that allowed us to utilize the intermediate traits as continuous variables.^27^ Although the linear mixed model approach controls to some extent the clonal population structure of *Mtb*, we discovered that the significant associations with RF under both carbon and antibiotic stress conditions were dominated by mutations shared by the L1.1.1.1 subclade of L1 strains (Fig. 3A, Fig. 3B, S5 Table).^40^ Indeed, the 32 *Mtb* clinical strains belonging to L1.1.1.1 drives the dramatic phenotypic differences between L1 with L2 and L4 (Fig. 2, fig. S7).

In our sample, the L1.1.1.1 clade shares 103 private mutations relative to all of the other strains (S6 Table). These variants include INDELs, nonsynonymous mutations, synonymous mutations, and intergenic mutations. With the L1.1.1.1 private mutations, we created a protein association network using the STRING database (Fig. 3C).^41^ For the largest interaction network, we performed a KEGG functional enrichment analysis (Fig. 3C, S6 Table, S7 Table), which indicated that mutations private to L1.1.1.1 are involved in fatty acid degradation (False Discovery Rate=0·00015), fatty acid metabolism (FDR=0·011), and propionate metabolism (FDR=0·046), amongst other pathways. These pathways track with the association with decreased fitness in the acetate + propionate condition (P=0·00036) (Fig. 3B, Fig. 3D).^42^

We also completed a SIFT analysis to assess which nonsynonymous mutations shared by L1.1.1.1 potentially affect protein function, as most likely candidates driving phenotype variation of the L1 subclade (Fig. 3C, S6 Table).^43^ Twenty of the 38 nonsynonymous mutations in proteins with matches within the SIFT database produce a premature stop codon or are a predicted intolerant mutation (Fig. 3C, S6 Table). This includes a premature stop codon in *mmpL5*, a gene encoding an efflux pump associated with BDQ resistance, which explains the GWAS link between L1.1.1.1 and BDQ susceptibility (P=1·26×10^-6^) (Fig. 3B-D).^44^ The L1.1.1.1 clade is also more susceptible to CFZ (Fig. 3A, fig. S7), resistance to which is also mediated by *mmpL5*.^44^ The GWAS did not return a statistically significant hit for PMD fitness, but L1.1.1.1 strains share a nonsynonymous mutation in *fgd1*, mutations in which are linked to PMD resistance.^45^

While the constellation of L1.1.1.1 mutations were the dominant feature of the intermediate phenotype GWAS, we did identify monogenetic associations with certain conditions, such as mutations in *nuoD*, which encodes a *Mtb* NADH dehydrogenase, linked to lower RF in acetate + propionate (P=4·59×10^-6^) (Fig. 3D); *Rv1339*, an enzyme involved in *Mtb* second messenger signaling, is associated with lower RF in glycerol (P=5·95×10^-6^) (Fig. 3B, Fig. 3D)^46^; and mutations in *Rv1707*, a putative sulfur transporter implicated in antibiotic persistence, is associated with increased RF in RIF (P=4·6×10^-6^) (Fig. 3B, Fig. 3D).^47, 48^ The latter was the only variant we discovered that was associated with increased fitness, suggesting that slow growth may provide *Mtb* with an adaptive advantage against stress.

### *Mtb* L1 clade is associated with poor clinical outcomes and increased transmission potential

We next sought to determine if the L1.1.1.1 phenogenomic determinants were associated with clinically relevant phenotypes. Interestingly, despite no previous association with the wider L1 lineage, we found that 44% of patients infected with strains from the L1.1.1.1 clade experienced cavitary disease, as opposed to 24% of patients infected by other strains in our sample (odds ratio = 2·49, confidence interval = 1·11-5·59, P=0·027) (Fig. 4A). In addition, 22% of patients infected with L1.1.1.1 strains failed treatment, compared to 6% of patients infected by other strains (odds ratio = 4·76, confidence interval = 1·53-14·78, P=0·0069) (Fig. 4A).

**Figure 4.**
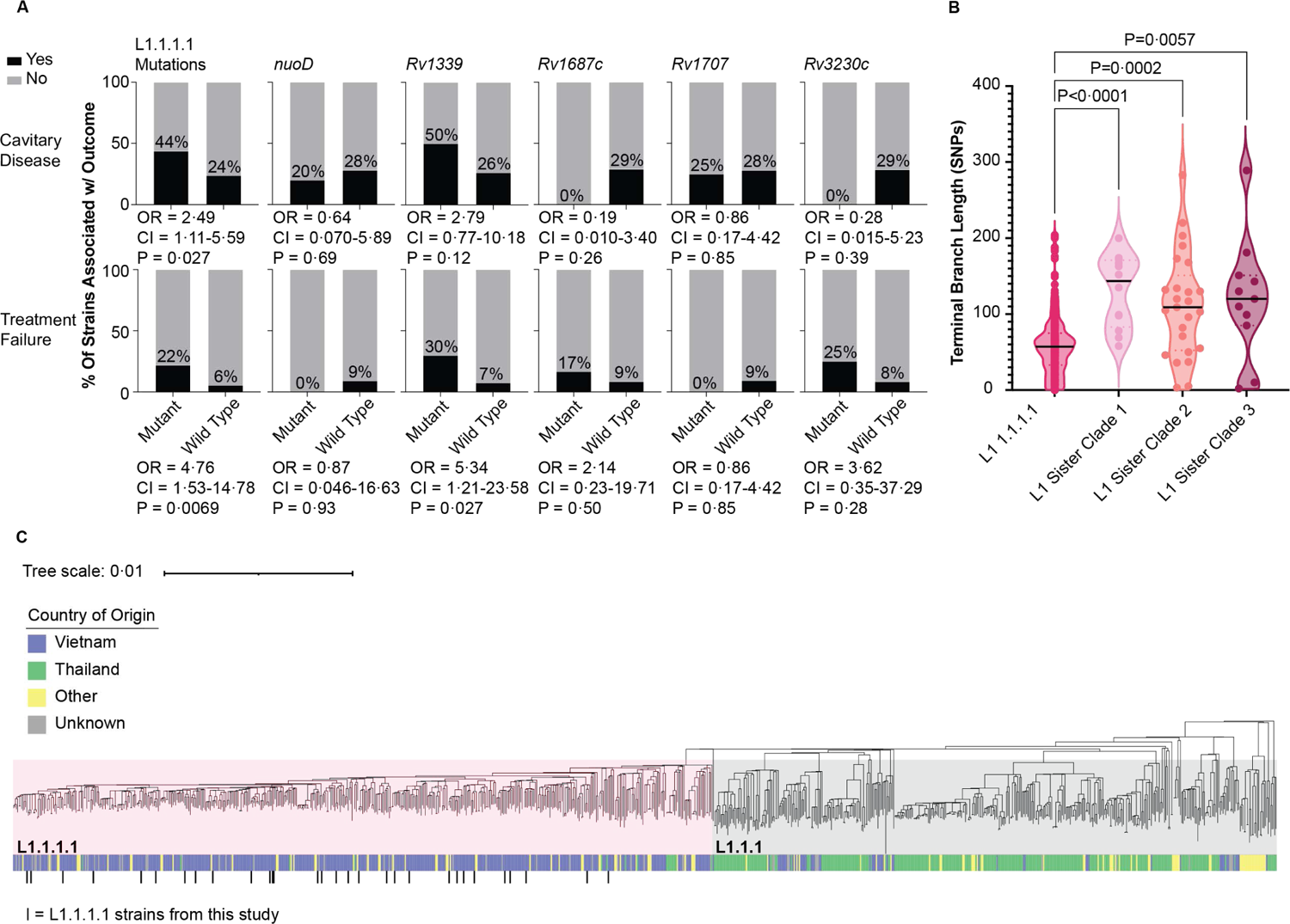
Lineage 1 clade associated poor patient outcomes and transmission in Vietnam. A. Percentage of strains associated with the indicated clinical outcomes, grouped by genotype for the indicated genes. Odd ratio, 95% confidence interval, and p-value shown. B. Distributions of terminal branch lengths determined by SNPs for different clades of L1. P-values from Kruskal-Wallis test and Dunn’s multiple comparison test in comparison to L1.1.1.1 indicated. C. Phylogenetic tree of 952 clinical isolates representing the global diversity of L1.1.1.1 and L1.1.1 *Mtb* clinical strains.

*Rv1339*, which has been implicated in modulating *Mtb* antibiotic susceptibility and is undergoing positive selection in TB patients globally was the only monogenic determinant associated with treatment failure (OR=5·34, CI=1·21-23·58, P=0·027), but not cavitary disease, suggesting disparate mechanisms may be accounting for these two distinct clinical outcomes associated with the L1.1.1.1 strains.^10, 46, 49^ We performed Mantel-Haenszel analyses to ensure that the associations we identified with L1.1.1.1 and *Rv1339* mutations are not explained by patient age or sex (S8 Table).

The finding that L1.1.1.1 strains have markedly lower RF under several *in vitro* conditions but were associated with worsened clinical outcomes might seem counter-intuitive. However, we have previously found that higher epidemiologic fitness of L2.2 strains correlates with slower growth in mice.^50^ Therefore, we sought to assess the epidemiologic fitness of the L1 subclade strains. Terminal branch lengths (TBLs), determined by the number of SNPs derived from the most recent branching point of each strain, serves as a proxy for the maximum evolutionary time after transmission.^11^ Using the entire panel of 1,635 clinical isolates in the transmission cohort, we compared L1.1.1.1 to three other closely related L1 clades.^11^ The TBLs of L1.1.1.1 strains are significantly smaller, which may be indicative of more recent transmission (Fig. 4B, fig. S8, S9 Table). In concert with the association we found between L1.1.1.1 strains and the clinical outcomes, this finding is consistent with the established relationship between cavitary disease, treatment failure, and transmission.^24, 25^

To assess the global relevance of our findings, we compared the L1.1.1.1 strains in our sample to a set of 952 clinical isolates that represent the global diversity of the L1.1.1.1 clade and the closely-related L1.1.1 subgroup (S10 Table).^10, 40^ Importantly, the L1.1.1.1 strains utilized in our analysis do not form a distinct monophyletic group within the L1.1.1.1 clade but are instead interspersed throughout (Fig. 4C). This suggests that the L1.1.1.1 intermediate phenotypes we describe may be widely representative of the entire clade. Consistent with previous observations, we find that 79·1% of L1.1.1.1 strains originated in Vietnam, while 69·4% of the L1.1.1 strains were isolated in Thailand.^11, 51, 52^ Therefore, these related clades demonstrate remarkable restriction within two distinct but proximal geographic regions. Accordingly, we show that L1.1.1.1 strains are phenotypically distinct from the remainder of L1 strains (fig. S7). Together, this data suggests that our characterization of the L1.1.1.1 intermediate traits are highly pertinent to the biogeographic context of Vietnam in particular and suggest the *Mtb* is following population-specific evolutionary trajectories.

## Discussion

Our work leverages a barcoded *Mtb* clinical collection to conduct high-throughput phenotypic screening across host and clinical stress conditions to identify lineage-level fitness differences across *Mtb* clades. The pipeline we created to barcode *Mtb* clinical strains to enable high-throughput competition experiments for phenotyping is conceptually similar to efforts to delineate determinates of *Plasmodium falciparum* antimalarial resistance and *M. abscessus* infection clearance.^53, 54^ However, in contrast to other studies with typical GWAS that try to identify single genetic determinates, we find that clade-level mutations with pleiotropic phenotypic effects may have a better predictive value for complex clinical phenotypes.

The most striking phenotype that we identified is slow growth under multiple conditions, which associated with a particular L1 subclade (L1.1.1.1) that was predictive of several clinically relevant outcomes— development of cavitary disease, likelihood of treatment failure, and possibly enhanced *Mtb* transmission. In addition to the L1.1.1.1 mutations, we also identified an association between *Rv1339* mutations across L1, decreased fitness in glycerol, and treatment failure. Consistent with our observation, *Rv1339* encodes a second messenger signaling enzyme that modulates drug susceptibility and is undergoing positive selections in clinical isolates.^10, 46, 49^ By identifying bacterial-intrinsic intermediate traits that link these mutations to the clinical phenotypes, we establish the plausibility that bacterial phenogenomic variation contributes to patient outcomes along with other factors with established associations with cavitary disease and treatment failure such as antibiotic adherence, comorbidities, drug resistance, and disease severity.^24, 55, 56\^

The relationship between decreased fitness in acetate + propionate, glycerol, and BDQ with the clinical and epidemiological outcomes is not immediately intuitive, as one might assume that increased fitness to stress would be a better predictor. However, *Mtb* is a slow-growing organism, which is hypothesized to be an adaption important for drug persistence and tolerance.^57^ Further, *Mtb* exhibits growth arrest in environmental conditions that simulate the immune-activated macrophage or granuloma, such as the combination of acidic pH and host-relevant carbon sources or low oxygen.^58, 59^ Therefore, decreased fitness in the stress conditions we observed could be an intermediate bacterial trait directly related to growth arrest and the mechanisms of the clinical outcomes. Alternatively, it is also possible that decreased fitness in conditions serve as markers of the processes directly driving the clinical outcomes.

The L1.1.1.1 subgroup carries clade-defining mutations in genes known to modulate resistance to not only BDQ but also CFZ and PMD, even though these mutations were acquired before adoption of these drugs in the antitubercular regimen.^60^ These data are consistent with other reports of subclade-defining mutations in *mmpL5* and *fgd1*, and our study provides further experimental evidence that these mutations alter *Mtb* clinical strain fitness to BDQ, CFZ, and PMD.^61–64^ This suggests that these strains have evolved mutations—likely in response to host pressures on bacterial energy metabolism or small molecule transport— which have caused altered sensitivity to these new antibiotics. These genomic changes in genes linked to altered BDQ, CFZ, and PMD sensitivity should be considered both to limit the emergence of additional drug resistance and in potentially tailoring TB treatment to maximize efficacy, especially considering the roll-out of the BPaL regimen – which includes BDQ and PMD.^60, 65^ This “precision medicine” approach could prove especially important for Vietnam, which has the highest burden of L1.1.1.1 infections.^11, 51, 52^ Indeed, the intermediate traits distinguishing the Vietnamese L1.1.1.1 strains could reflect the phenotypic effects of bacterial adaption to unique host and environmental pressures in this region. Future should work is needed to define the mechanisms driving biogeographical restriction of this clade. Further, our TBL analysis may indicate that this clade is undergoing a recent expansion event in Vietnam. Because TBLs may be shaped by extenuating factors -- such as bacterial mutation rate, the genomic cluster rate, sample size, and population size -- additional analyses that are outside of the scope of this study must be employed to assess transmission dynamics more definitively. While the field has largely focused on the global pandemic success of L2.2, we provide experimental evidence highlighting the clinical significance of the geographically-restricted L1.1.1.1 subgroup. Increased transmission within Vietnam should motivate follow-up concerning the treatment outcomes and altered drug sensitivities that we identified.

The diversity of intermediate phenotypes displayed by the *Mtb* clinical strains is intriguing because *Mtb* was historically considered to be a genetically monomorphic organism.^66^ The field recognizes that *Mtb* clinical outcomes are heterogenous, and our data suggests that bacterial heterogeneity should not be overlooked when considering factors contributing to variation in patient TB treatment and disease phenotypes.^67^ Importantly, this heterogeneity, while having a phylogenetic relationship, is not easily distinguished at the canonical *Mtb* lineage level. Given the genomic diversity within *Mtb* lineages, it may be that other sub-lineages can be associated with similar negative clinical outcomes but rely on an alternative distinct combination of genomic mutations and manifest with different phenotypes. However, it would be interesting to assess whether the slow growth intermediate phenotypes we observed across multiple stress conditions is relevant in different phylogenetic contexts.

Together, our work highlights the utility of conducting high-throughput phenotyping of clinical strains to define intermediate fitness phenotypes associated with negative clinical outcomes and leverage these phenotypes to discover sub-lineages (and genomic mutations) of interest. These phenotypic and genomic features may prove useful as both tools for dissecting mechanisms of *Mtb* pathogenesis as well as clinically relevant biomarkers associated with treatment failure.

