## Supplementary Figures for "High-throughput phenogenotyping *of Mycobacteria tuberculosis* clinical strains reveals bacterial determinants of treatment outcomes"

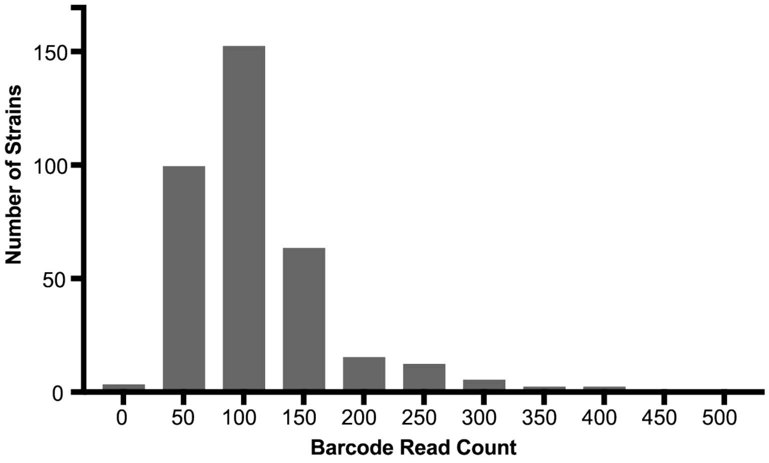


**Supplementary Figure 1.** Histogram of the barcode read counts in the barcoded input library of clinical strains. Average of three technical replicates. 159 unique strains with 1-3 different barcodes each, n = 355. Most strains (315 of 355) have barcode read counts between 50-150.

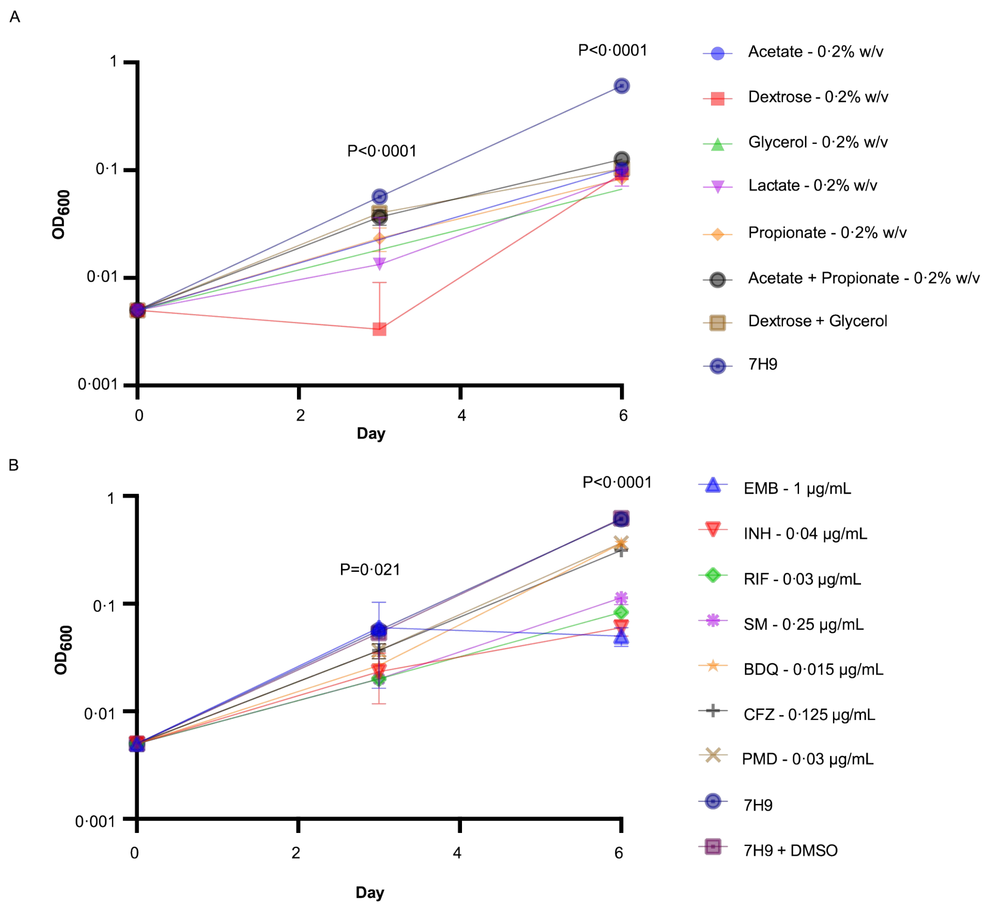


**Supplementary Figure 2.** Growth curves of the barcoded *Mtb* clinical strain library days 3 and 6 after inoculation with the barcoded input library of clinical strains. All cultures started at $OD_{600}$ 0.005 at day 0. Triplicate replicates shown. Dots indicate mean, error bars indicate standard deviation. P-value indicates results from ordinary one-way ANOVA for each timepoint. A. For day 3 relative to 7H9, Dunnett's multiple test correction p-value is <0·0001 for acetate, dextrose, and glycerol; p-value = 0·002 and 0·0022 for lactate and propionate respectively; not significant for acetate + propionate and dextrose + glycerol (p-value = 0·079 and 0·18 respectively). For day 6, all conditions compared to 7H9 has p-value <0·0001. B. For day 6, the Sidak's multiple test correction p-value is >0·99 for EMB, 0·12 for INH, and 0·069 for SM, all relative to 7H9. Relative to 7H9+DMSO, the p-value is 0·12 for RIF, 0·80 for CFZ, 0·31 for BDQ, and 0·80 for PMD. For D6, P<0·0001 for all conditions.

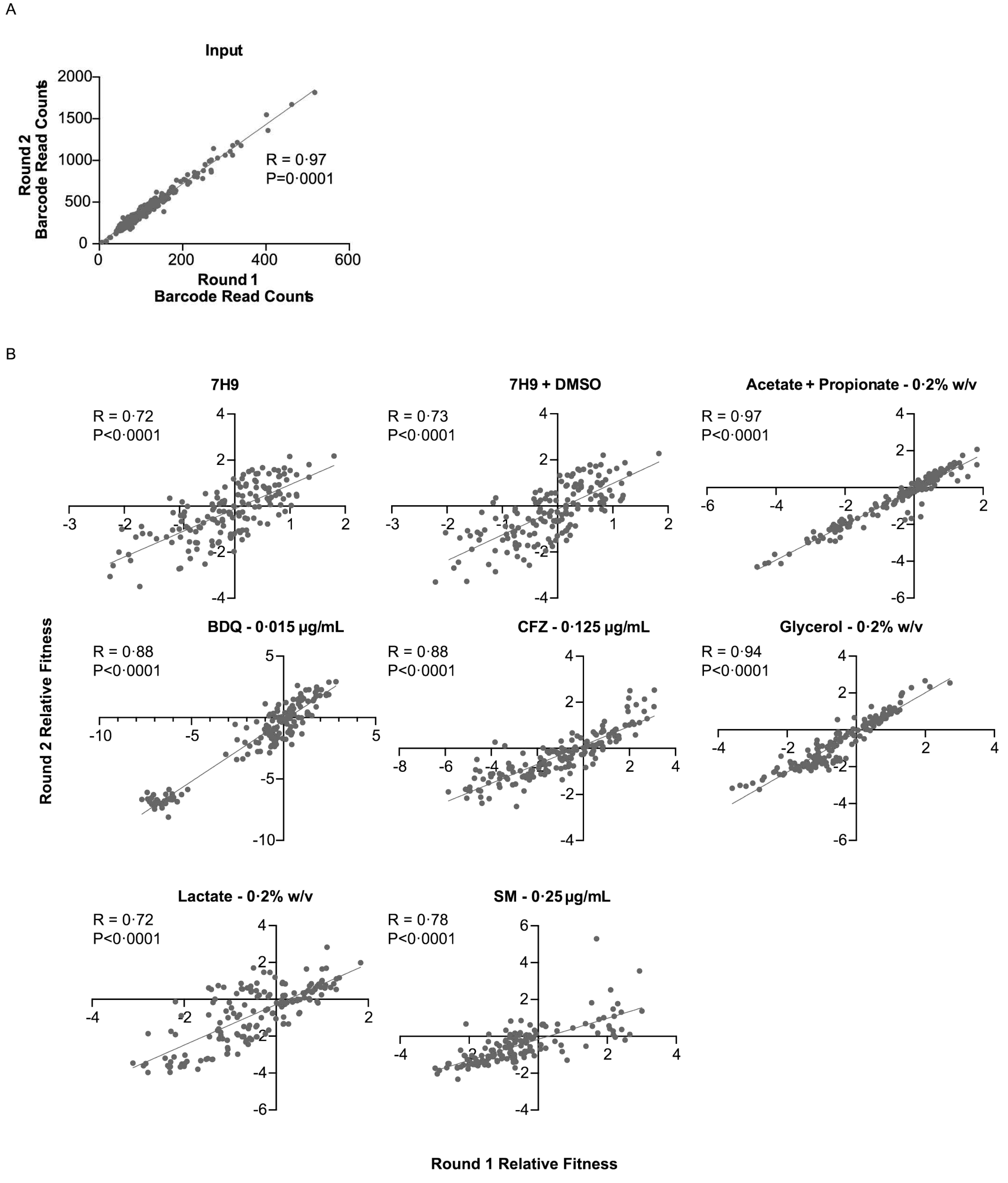


**Supplementary Figure 3.** Correlation of barcode read counts (A) or RF values (B) between two independent rounds of competition experiments. Spearman correlation coefficient indicated.

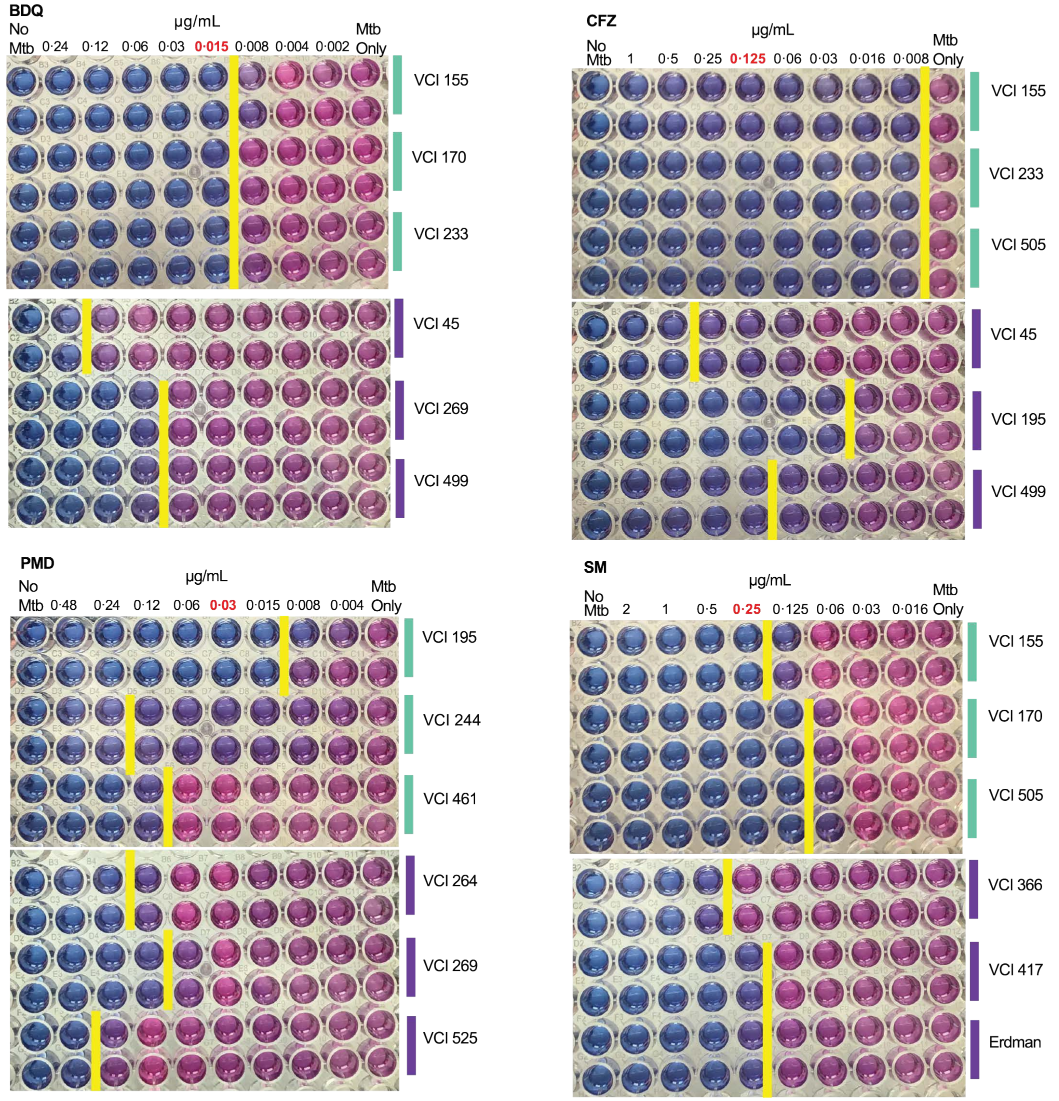


**Supplementary Figure 4.** Alamar blue MIC assays for 3 strains among the most fit and least fit as listed in S4 Table. Completed in duplicate. The concentrations used in the competition experiments is in red. MIC is the concentration to the left of the yellow bar.

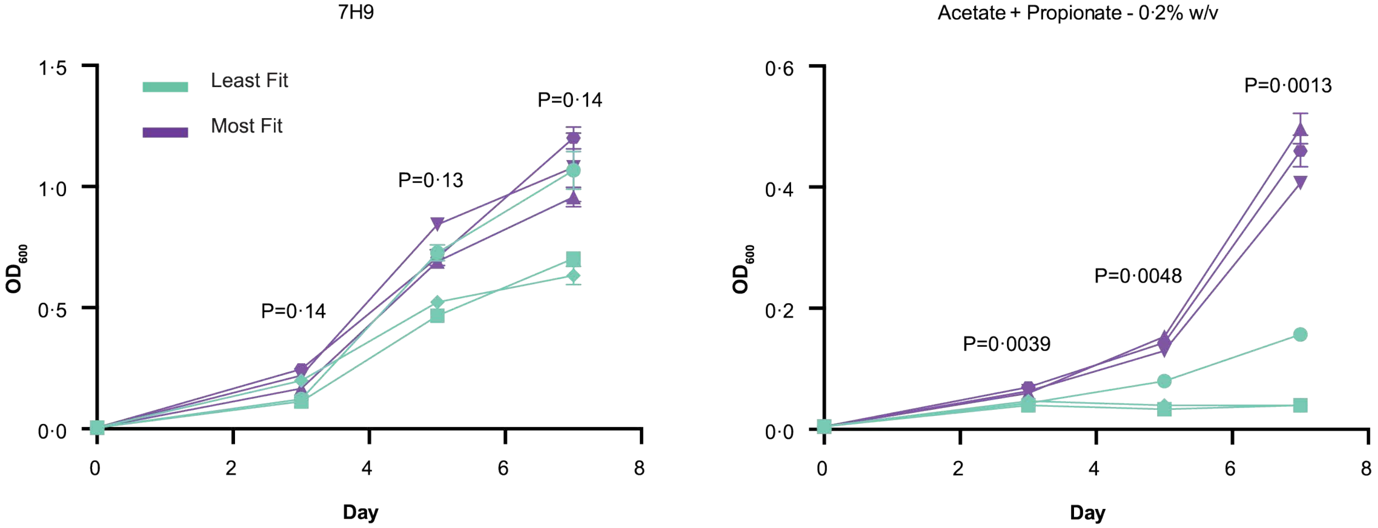


**Supplementary Figure 5.** Growth curves of strains among the most fit and least fit in the acetate + propionate competition experiment. Completed in triplicate for each strain. Error bars indicate standard deviation. Results of an unpaired t-test are shown. Strains tested listed in S4 Table.

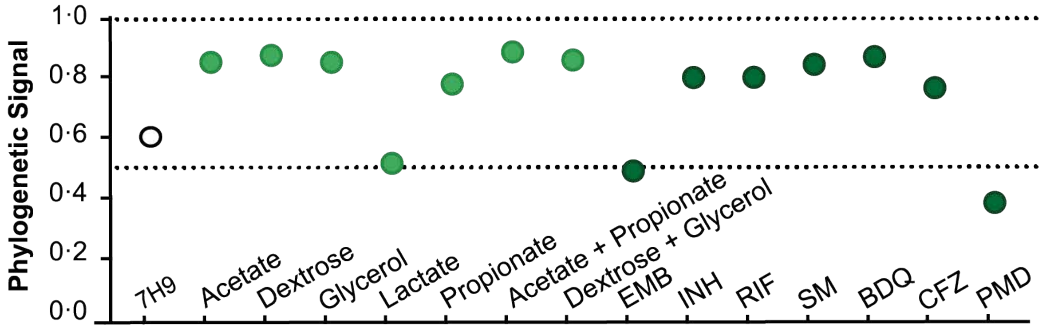


**Supplementary Figure 6.** Phylogenetic signal (lambda) for each condition based on D6 RF values. All p-values <0·0001.

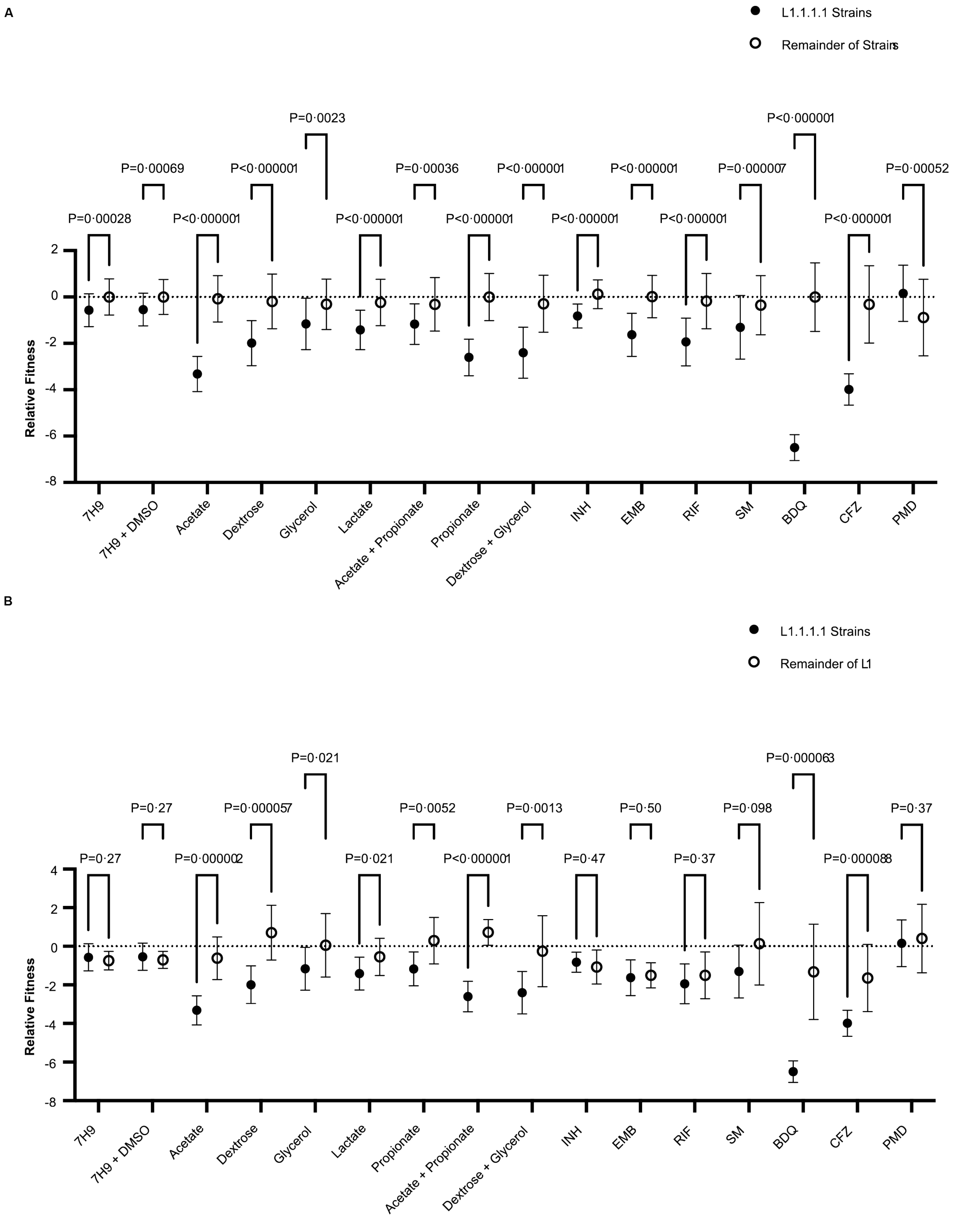


**Supplementary Figure 7.** Relative fitness values of the L1.1.1.1 strains compared to the remainder of strains (A) and remainder of L1 strains (B) across all conditions, p-value results of Mann-Whitney test shown after Benjamini, Krieger, and Yekutieli multiple test correction. Dots indicate average, error bars indicate standard deviation.

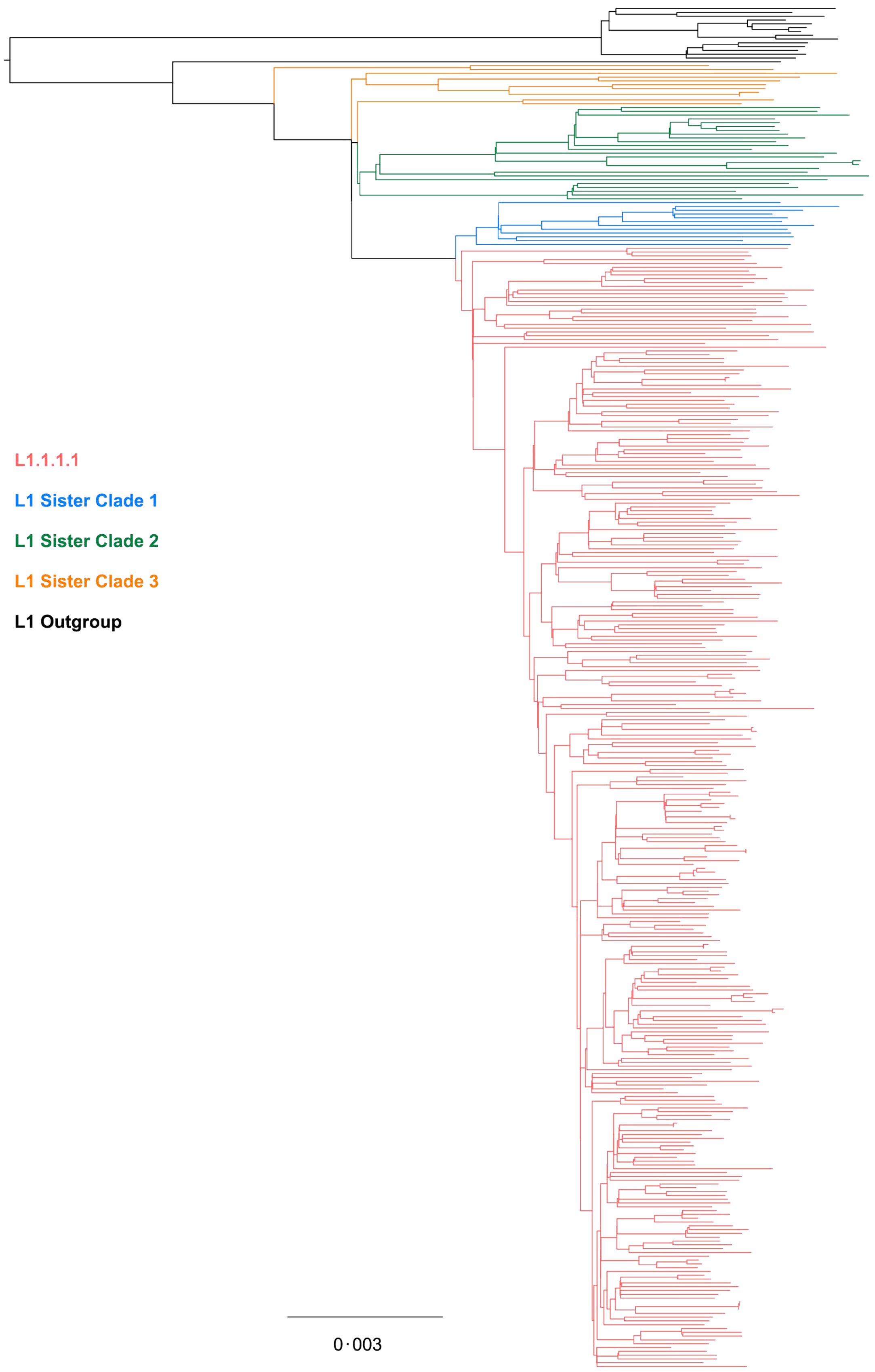


**Supplementary Figure 8.** Approximately maximum-likelihood tree of the 358 L1 strains analyzed for the terminal branch length analysis, colored by relationship to L1.1.1.1. Scale indicates the number of mutations per site. Visualized with FigTree v1.4.4.
