## Extended Methods for "High-throughput phenogenotyping *of Mycobacteria tuberculosis* clinical strains reveals bacterial determinants of treatment outcomes"

**Mtb clinical strains**

The strains were originally isolated from patient sputum samples cultured on Lowenstein-Jensen slants for 4-6 weeks at 37° C then cryo-archived at -20° C in 7H9 media (7H9 Middlebrook salts, 0·2% glycerol, 10% OADC, 0·05% Tween80) with 40% glycerol. In preparation for shipping to our facilities, the isolates were transferred to 50 mL tubes containing 15 mL of 7H9 media and incubated at 37° C for 2 months without shaking. The cultures were then centrifuged, and the supernatant was removed before 1 mL of 7H9 media with 40% glycerol was added. The content was mixed well before transfer into 2 mL external threaded screw-capped tube with label. The cap was covered with parafilm and stored at -20° C before shipping. Upon receipt of the cultures, we inoculated the strains into inkwell containing 10 mL of 7H9 media and incubated the strains at 37° C with shaking until the cultures became turbid. The strains were then stored at -80° C in 7H9 media with 25% glycerol.

**Drug sensitivity testing**

To confirm the samples used in our study were phenotypically sensitive to rifampicin (RIF) before further processing, we inoculated each isolate in an inkwell with 10 mL of 7H9 media and incubated at 37° C with shaking until the strains reached an OD_600_ of 0·6-0·8 as measured by a spectrophotometer. The strains were then diluted to an OD_600_ of 0·05 and pooled together by adding 2 mL of each strain to a 100 mL inkwell containing 7H9 media and 1 μg/mL of RIF. Each pool contained 2-15 different strains. The OD_600_ of the pools was measured on days 1, 3, 5, 7, 10, and 14 of growth at 37° C with shaking. No growth was detected in the pools comprising the strains from this study. Sensitivity to RIF was further determined by plating 100 uL of the OD_600_ 0.05 diluted pool on 7H10 (7H10 Middlebrook salts, 0·2% glycerol, 10% OADC) with 1 μg/mL of RIF. The plates were checked for lack of colonies after 3 weeks of growth at 37° C.

**Barcoded Mtb clinical strain library**

The barcoding plasmids were cloned by digesting the plasmid pJeb402 with the restriction enzymes Kpn1 and MluI and ligating with gBlocks (IDT, Coralville, Iowa, USA) to form the barcodes which integrate at the L5 phage site. The plasmid contained qTag-29 from Blumenthal *et al*.^1^ For biological replicates, 1-3 colonies from each of the 159 strains were selected containing unique barcodes as confirmed by Sanger sequencing amplicons produced with primers TTTACGGTTCCTGGCCTTTTGC and GGTCTCCCCATGCGAGAGTAGG.

To create a pooled barcoded library, the 355 barcoded strains were individually inoculated in 10 mL of 7H9 media in inkwells with 20 μg/mL kanamycin and incubated at 37° C with shaking until they reached OD_600_ of ~0·6. The strains were diluted to an OD_600_ of 0·5 and 1 mL of each diluted cultured was combined to create the barcoded pool. We aliquoted the pool into 2 mL freezer stocks for subsequent experiments.

***In vitro* competition experiments**

For the competition experiments, the antibiotics were added at the indicated concentration to 10 mL of 7H9 with 20 μg/mL of kanamycin in inkwells. The carbon source conditions were set up by adding the indicated concentration of carbon source to 10 mL of 7H12 media (7H9 Middlebrook salts, 0·1% casamino acids, 0·05% tyloxapol) with 20 μg/mL of kanamycin in inkwells. For conditions containing two carbon sources, 0·1% w/v of each carbon sources was utilized. The pooled clinical strain library (input library) was inoculated into each condition to an OD_600_ of 0·005. Six inkwells were prepared for each condition to serve as technical replicates. The cultures were incubated at 37° C with shaking and the OD_600_ was measured on D3 and D6 to monitor bulk library growth. At each time point, the gDNA was extracted from the entire biomass for three technical replicates per condition. The gDNA extraction was performed according a previously described protocol.^2^ During this step, a D6 lactate technical replicate and some D3 technical replicates for acetate, dextrose, propionate, acetate + propionate, dextrose + glycerol, and EMB were lost. The gDNA from the input library was also extracted in triplicate. We then prepared libraries for MiSeq V3 sequencing (Illumina, San Diego, CA, USA) using custom primers that amplify the barcode.^2^

Random sequences that serve as unique molecular counters were added to the amplicons during library prep to control for barcode application bias.^3^ The FASTQ files generated from sequencing were searched for a unique molecular counter and a strain-specific barcode. For each sample for each read, the added unique molecular counter was identified using the following regular expression ("C([ACTG]{3})C([ACTG]{3})C([ACTG]{3})GCGCAACGC") and the barcode was counted using exact string matching from a list of barcode sequences with a custom python script. The barcode sequences and list of FASTQ files to be analyzed were supplied as a text file and read into the script. Total read counts for each barcode and the unique counts are output as separate files. The raw barcode read counts were then adjusted based on the unique molecular identifier (UMI) to calculate strain abundance.

**Analysis of competition experiment barcode sequencing data**

To analyze the technical variation from this experiment, the UMI-adjusted read counts for each individual barcode was normalized to the total reads counts in a given technical replicate to control for differences in read depth across the samples. For each growth condition, every barcode was then normalized to the corresponding input library value averaged across the input library technical replicates (S2 Table). This step controlled for differences in strain abundances in the input library. A replicate of EMB was removed due to low sequencing depth (<1000 reads in the sample). A principal component analysis to compare technical replicates was performed using these values with the stats package (version 4.3) of R (version 4.2.2).

In order to analyze the biological replicates (identical strains with unique barcodes), the input normalized values for each barcode were averaged across technical replicates for each condition (S2 Table). These averaged normalized read counts were utilized to complete the Pearson correlation coefficient analysis where we compared the correlation between every unique barcode (n=355) across all conditions. For the biological replicates, comparisons of the averaged input normalized read counts that resulted in a Pearson correlation coefficient of 0·62 or less was deemed an outlier via the ROUT test, Q = 1%, performed in GraphPad Prism V9 (GraphPad Software, San Diego, CA, USA) based on the distribution of all of the biological replicate R^2^ values.^4^ As a result, 32 of 355 barcodes were removed from downstream analysis.

To obtain the relative fitness values for each strain, we averaged the input normalized read counts for the technical and biological replicates together and found the log_2_ of these values after removing the outlier barcodes (S2 Table). The relative fitness values were utilized for the D3 vs D6 dot plots, AGNES hierarchal clustering analysis (cluster package of R, version 2.1.4), the phylogenetic signal analysis (phytools package of R, version 1.2.4), and the GWAS.^5^

For the D3 vs D6 dot plots, the D3 samples from the first round of competition experiments were analyzed as described above to generate relative fitness values (S3 Table). The Spearman correlations comparing the two independent rounds of competition experiments were generated by determining the relative fitness values for the second round of competition experiments also as described above (S3 Table).

**Genomic, phylogenetic, and mutation analyses**

The gDNA extraction of the clinical strains for WGS was completed according to previously published protocols.^2^ We also whole-genome sequenced the Erdman strain we utilized. Alignment, variant calling, and mutation annotation were performed as described by Hicks *et al* to create a joint variant call file (VCF) with the following modifications: reads were aligned to the inferred ancestral genome of the *Mycobacterium tuberculosis* complex most recent common ancestor; read duplication, coverage, depth, and also insert size were assessed with Picard tools (version 2.25.7) of Java (version 1.8); and SNPs and small INDELs were determined using the HaplotypeCaller tool from the Genome Analysis Toolkit (version 4.3).^3,6^ Sites with a mapping quality score of less than 50 were removed from the VCF using VcfFilter (version v0.2), along with sites within low confidence or repetitive regions using VCFtools (version 0.1.14).^7–9^ We constructed a whole-genome phylogenetic tree based on 33,566 sites from the joint VCF using fastTree according to the method described by Hicks *et al.*^3,10^ A *Mycobacterium canettii* strain (GenBank accession NC_015848) served as the outgroup. The phylogenetic tree was visualized with iTOL (version 6.6).^11^ *M. canettii* was removed from the tree before rooting at the midpoint. Holt *et al* completed lineage typing of the strains, which we refined based on the phylogenetic reconstruction.^12^ We based our L1.1.1.1 typing according to the system described by Coll *et al*.^13^

The GWAS was performed using the linear mixed model option of the Python package (Version 3.8.5) of Pyseer (version 1.3.10).^14^ The joint VCF described above provided the polymorphisms after removal of synonymous sites, INDELS, and the *M. canettii* sequence. Sites with multiple alleles were formatted for Pyseer compatibility using a custom script. In addition, the VCF was normalized and the file compressed and indexed using BCFtools (version 1.5). Relative fitness values generated from the competition experiments were treated as continuous phenotypes. The phylogenic tree described above was used to create a distance and similarity matrix to control for population structure. We included the lineage assignment for each strain with the lineage clusters option to control for lineage effects. We utilized the burden test option to group sites according to genes and intergenic regions. To determine the significance threshold of 2·13x10^-5^, we performed Bonferroni correction based on the number of unique variant patterns and a significance level of 0·05.

The protein association network of the L1.1.1.1 private mutations was created via the Cytoscape software (version 3.9.1).^15^

For sfig. 8 and Fig. 4C involving the terminal branch length analysis and the global L1.1.1.1 comparison, the fixed SNP calling and phylogenetic reconstruction was performed as described by Liu *et al*.^16^
